# A study on edge devices for image classification of the Tasmanian devil (*Sarcophilus harrisii*) for vaccine delivery

**DOI:** 10.1101/2025.09.29.679120

**Authors:** Prithul Chaturvedi, Andrew S. Flies, Soyeon Caren Han, William M. Connelly

## Abstract

1. A target-specific bait dispenser is required for oral bait vaccination of the endangered Tasmanian devil (*Sarcophilus harrisii*) against the deadly devil facial tumour disease. Development of a broadly extendible dispenser would be a beneficial ecological tool. A camera-based edge device with an onboard deep learning model can be used to identify the target species using image classification. It is important to choose a suitable edge device for the dispenser’s effective application in power constrained environments.
2. We evaluated four edge devices for the smart bait dispenser-ArduinoPro Nicla Vision, ArduinoPro Portenta H7 with Vision Shield (LoRa), Raspberry Pi 3 Model B+ and Raspberry Pi Zero 2 W. Two simple convolutional neural networks, and four fine-tuned pretrained models (MobileNetV2, MobileNetV3Small, ResNet50V2, and ResNet152V2) were trained on trail camera images of devil and non-devil species. These models were evaluated across four metrics and post-training quantised for deployment. The edge devices were assessed on inference latency of each model in seconds (s) and power consumption in watt (W).
3. We found that a simple CNN based image classification model yielded the best overall result of ∼96% across all metrics. Raspberry Pi 3 Model B+ and Zero 2 W could run all the six models whereas Portenta failed to run the two ResNet models and Nicla Vision failed to run all four pretrained models. All edge devices were found to be quick enough for dispenser application (average inference latency of 0.718 s). Portenta consumed the lowest power during inference (0.19 W), idle (0.185 W), and light sleep (0.045 W) states whereas Nicla Vision consumed the lowest power during deep sleep (0.002 W).
4. Overall, we found that simple traditional CNNs was best suited for species classification in a camera-based smart bait dispenser for vaccination of Tasmanian devils. We conclude that ArduinoPro Portenta H7 with Vision Shield is the most competent device for this edge AI application. We suggest integration of this system with a microcontroller unit like ATtiny85 to minimise inactive power consumption of the dispenser. This method can be readily adapted for other species in vaccination and supplemental feeding projects.

## 1 INTRODUCTION

The Tasmanian devil (*Sarcophilus harrisii*; hereafter devil) is the largest extant carnivorous marsupial and the native apex predator of Tasmania, Australia. The species has been listed as endangered by the International Union for Conservation of Nature and Natural Resources (IUCN) since 2008 (Hawkins, 2008), largely due to the emergence of two transmissible cancers-devil facial tumour 1 (DFT1) and DFT2. Each of the tumours can independently cause devil facial tumour disease (DFTD). The disease has caused regional declines of up to 82% with nearly 100% case fatality rates (Cunningham et al., 2021).

Oral bait vaccines (OBVs) have been widely used as a wildlife disease management strategy for rabies (reviewed in (Rupprecht et al., 2024)). A vaccine against DFTD that could be delivered in edible baits would be a powerful management and conservation tool (Flies et al., 2020). These baits can be dropped from aircrafts or hand distributed in target population areas. Whilst these approaches are effective at a broad scale, they can be costly due to consumption by non-target species and consumption of multiple baits by a single individual. Maximising bait uptake by devils while minimising uptake by sympatric species would be a key for economical and effective control of DFTD.

In ecological systems where non-target competition is a significant challenge, bait dispensers have been used (Bastille-Rousseau et al., 2024; Smyser et al., 2015; Whisson & Salmon, 2009). Initial trials of ground distribution of placebo baits in Tasmania resulted in devils consuming around 7% of the baits; eastern quolls (*Dasyurus viverrinus*), brushtail possums (*Trichosurus vulpecula*), and Tasmanian pademelons (*Thylogale billardierii*) consumed approximately 75% of the baits (Dempsey et al., 2022). Use of a timer-based bait dispenser developed for raccoons (*Procyon lotor*) in North America (Smyser et al., 2015) was trialled in Tasmania and resulted in a nearly tenfold increase in bait uptake by devils (Dempsey et al., 2022). However, eastern quolls often removed the baits from the dispenser before devils arrived. Furthermore, the relatively small home range of eastern quolls led to individuals frequently returning to the dispenser to retrieve additional baits

A dispenser that implemented species recognition technology could reduce bait consumption by non-target species and better limit the number of baits consumed by an individual. Designing a dispenser that could be rapidly adapted to target alternative species would be a broadly useful conservation tool. This would have to balance multifunctionality with the increased cost of adding features and potential reductions in durability.

Deep learning (DL) is a sub-field of machine learning (ML) that extracts higher-level features from data (e.g. images) with limited feature engineering (Choudhary et al., 2022; LeCun et al., 2015). DL methods are applicable in a wide range of domains due to its scalability, generalisability, and robustness (Alzubaidi et al., 2021). However, they are computationally demanding and are typically deployed over high-performance infrastructures or distributed clusters in data centres (Guo, 2018). Here, input data is sent to another location for processing via a network. This requires a secure and private connection for deployment, which may not be feasible in field applications. Alternatively, it is possible to run scaled-down DL models on smaller and cheaper edge devices like single board computers (SBCs) and microcontroller unit (MCU) boards (reviewed in (Kong et al., 2022)). Here, input data are captured and processed in the same location, enabling secure real-time processing (Warden & Situnayake, 2019) and allowing decisions to be made in the field. SBCs and MCU boards, coupled with a camera, are used to deploy DL methods such as object detection and image classification (reviewed in (Iqbal et al., 2024) and (Maheepala et al., 2021)). This is also referred to as edge artificial intelligence (edge AI).

Edge AI applications in camera traps and acoustics have been used in conservation for species recognition to remotely monitor wildlife populations and to detect poaching and illegal deforestation (Fergus et al., 2024; Schwartz et al., 2021). A five-bait dispenser prototype was successful in targeting feral dogs (*Canis familiaris*) based on real-time camera trap images using object detection method (Charlton et al., 2023). This system is based on Raspberry Pi 3 Model B+ that requires a large battery to provide power for multiday field applications, making execution costly and setup bulky. Raspberry Pi Zero 2 W is a smaller and more power efficient alternative but has comparatively limited computing power. Alternatively, using a high-end microcontroller unit device that offers comparable computational capabilities for image-based species classification can run the dispenser longer on smaller batteries. Overall, balancing edge device capabilities with the power requirements is crucial for field applications.

In this study, we compare four edge devices for a smart bait dispenser-two Arduino-based microcontroller unit devices, ArduinoPro Nicla Vision and ArduinoPro Portenta H7 with Vision Shield (LoRa), and two Raspberry Pi-based single board computer devices, Raspberry Pi 3 Model B+, and Raspberry Pi Zero 2 W. We conducted a preliminary assessment of 31 convolutional neural network (CNN) models to identify the most suitable models for devil classification on edge devices. Here, we discuss six of the best performing models based on accuracy, precision, recall, and F1-score metrics. The edge devices were scrutinised on inference latency and power consumption, as these are the most important factors for successfully implementing the bait dispenser in energy deficit remote locations.

## 2 METHODS

### 2.1 Dataset

The images used in this study were provided by the Department of Natural Resources and Environment Tasmania. These 1920×1080 pixels photos were collected during routine wildlife monitoring activities in Tasmania between 2015 and 2020 using Reconyx HC500 trail cameras. The dataset contained images of relevant species: devil, eastern quoll, spotted tail quoll (*Dasyurus maculatus*) wombat (*Vombatus ursinus*), brushtail possum (*Trichosurus vulpecula*), Bennett’s wallaby (*Notamacropus rufogriseus*), Tasmanian pademelon, feral cat (*Felis catus*), bandicoots (*Isoodon obesulus* and *Perameles gunnii* species), unidentified small mammals like rodent (presumable *Rattus spp.* and *Mus spp.*), snakes, cattle, birds, and empty background (Figure 1).

**Figure 1.**
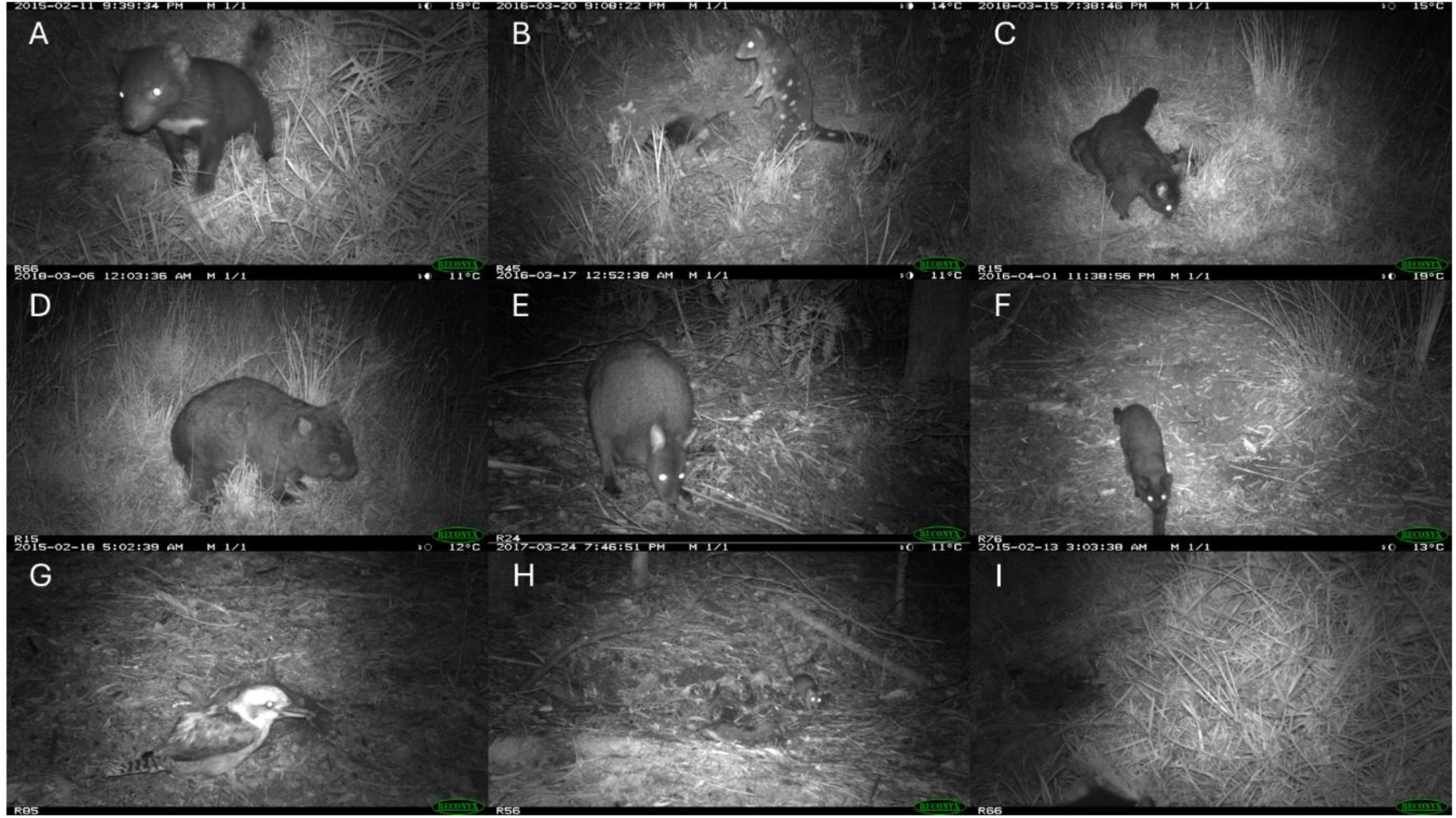
Example images used in training image classification models. (A) Tasmanian devil, (B) spotted tail quoll, (C) brushtail possum, (D) wombat, (E) wallaby, (F) feral cat, (G) kookaburra, (H) rodent, and (I) background.

A total of 9315 images were manually annotated as devil and non-devil. The devil class consisted of images where clearly distinctive features of a Tasmanian devil were visible face (eyes, ears, hairless snout, DFTD tumour, mouth/teeth), tail, and white markings on chest, arm, or back. During initial experiments, we found that partial images of wombats and possums generated false-positive predictions across all models. To minimise this, we created a “devil without distinctive features” subclass under non-devil class. The other subclasses under non-devil class were cat, wombat, macropods, quolls, possum, and miscellaneous. The miscellaneous class was created to include images of species like rodent, bandicoot, birds, and cattle that are seldom caught on trail cameras, and empty background.

The dataset was randomly split into training (80%, including validation) and testing (20%) subsets. To ensure that our data contained a representative distribution of species, stratified randomisation was used on the non-devil class. To improve intra-class variability, each image was passed through a ‘data augmentation layer’ consisting of random horizontal flips and random rotation at factor = 0.05 before feeding into models. The images were rescaled to 54×54 pixels for the classification algorithms to meet the memory limitations of the MCUs.

### 2.2 Model training

All computations were carried out using Python 3 on JupyterLab hosted on the Australian Research Data Commons’ Nectar Research Cloud. Keras application programming interface was used in model development (Chollet, 2015).

#### 2.2.1 Models

For this study, we considered 16 custom trained convolutional neural networks (CNN) and 15 pretrained models for transfer learning. This was to experimentally identify the most suitable candidates for binary image classification on edge devices. Here we focus on the best two CNNs and the best four pretrained models. The architectures of the chosen CNNs are in Table 1. The top performing pretrained models were ResNet50V2, ResNet152V2 (both architectures in (He et al., 2016)), MobileNetV2, MobileNetV3 (architectures in (Howard, 2017) and (Howard et al., 2019), respectively).

**Table 1.** Architectures of models (A) CNN1 and (B) CNN2.

**A. Model CNN1**
| Layer Name | Type | Output shape | Parameters |
| --- | --- | --- | --- |
| 1.1 | Conv2D | (54, 54, 32) | 896 |
| 1.2 | MaxPooling2D | (27, 27, 32) | 0 |
| 2.1 | Conv2D | (27, 27, 32) | 9,248 |
| 2.2 | MaxPooling2D | (13, 13, 32) | 0 |
| 3.1 | Conv2D | (13, 13, 16) | 2,064 |
| 3.2 | MaxPooling2D | (6, 6, 16) | 0 |
| 4 | Dropout (0.25) | (6, 6, 16) | 0 |
| 5 | Flatten | (576) | 0 |
| 6 | Dense | (1) | 577 |

**B. Model CNN2**
| Layer Name | Type | Output Shape | Parameters |
| --- | --- | --- | --- |
| 1.1 | Conv2D | (54, 54, 32) | 896 |
| 1.2 | MaxPooling2D | (27, 27, 32) | 0 |
| 2.1 | Conv2D | (27, 27, 32) | 9,248 |
| 2.2 | MaxPooling2D | (13, 13, 32) | 0 |
| 3.1 | Conv2D | (13, 13, 16) | 2,064 |
| 3.2 | MaxPooling2D | (6, 6, 16) | 0 |
| 4.1 | Conv2D | (6, 6, 16) | 1,040 |
| 4.2 | MaxPooling2D | (3, 3, 16) | 0 |
| 5 | Dropout (0.25) | (3, 3, 16) | 0 |
| 6 | Flatten | (144) | 0 |
| 7 | Dense | (1) | 145 |

Total parameters: 13,393

#### 2.2.2 Model training hyperparameters

a. Image size: 54×54 (dimensions) ×3 (channels) pixels
b. Batch size: 128 images
c. Loss function: Binary cross-entropy
d. Optimiser: Adaptive Moment Estimation (Adam)
e. Learning rate:

i For CNNs: 10^-3^
ii For pretrained models: 10^-3^ for training the top layer and 10^-5^ for fine tuning
f. Epochs:

i For CNNs: 300
ii For pretrained models: 50 for training the top layer and 250 for fine-tuning

#### 2.2.3 Post-training quantisation

All full-size models were quantised to reduce model size and, consequently, latency on edge devices at the cost of an acceptable dip in model performance using the following three techniques:

1. Dynamic range Float32 (FP32) quantisation: Model weights and activations are converted to 8-bit integers (INT8), shrinking to approximately one quarter of its original size; integers are dequantized back to FP32 values during inference. F32 models can only run on SBCs.
2. Float16 (F16) quantisation: Model weights and activations are converted to F16 values, shrinking model size to half. F16 models can only run on SBCs.
3. Full integer quantisation: Model weights and activations are converted to INT8, shrinking model size to quarter. INT8 models can run on both MCUs and SBCs.

The intent behind exploring three types of post-training quantisation methods was to thoroughly assess the performance metrics of full models against smaller counterparts for edge devices. Since our goal is to evaluate two SBC and two MCU, running selected I8Q models provides head-to-head comparison. TensorFlow Lite (also known as LiteRT) was used for post-training quantisation (Google AI for Developers, 2025).

#### 2.2.4. Model evaluation

The optimal classification threshold for each original and quantised model was determined by the maximum Youden’s J statistic on the receiver operating characteristic (ROC) curve. If fpr and tpr are false positive and true positive rates respectively, then the optimal threshold t is characterised by:

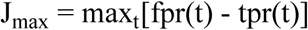

We chose accuracy, precision, recall, and F_1_-score as the metrics to assess each model’s predictive performance on the test dataset. If F_P_, F_N_, T_P_, and T_N_ denote false positive, false negative, true positive, and true negative respectively, then:

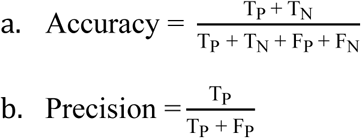

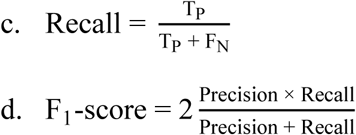

Scikit-learn was used to evaluate all full-size and quantised models (Pedregosa et al., 2011).

### 2.3 Edge devices

The following devices were considered for use in the smart dispenser in this study:

1. ArduinoPro Nicla Vision (hereafter Nicla Vision) with GC2145 onboard camera (Arduino, 2022; GalaxyCore, 2013).
2. ArduinoPro Portenta H7 with Vision Shield (LoRa) (hereafter Portenta) with HM01B0 onboard camera (Arduino, 2020; Himax, 2016).
3. Raspberry Pi 3 Model B+ (hereafter RPi 3B+) with Waveshare OV5647 camera module (Raspberry Pi, n.d.; Sparkfun, 2009; Waveshare, n.d.).
4. Raspberry Pi Zero 2 W (hereafter RPi Z2W) with Waveshare OV5647 camera module (Raspberry Pi, n.d.; Sparkfun, 2009; Waveshare, n.d.).

#### 2.3.1 Firmware/operating system

OpenMV firmware v4.7 was used for microcontroller boards Nicla Vision and Portenta. Raspberry Pi OS Legacy 32-bit Lite Debian v11 Bullseye was used for single board computers RPi 3B+ and RPi Z2W.

#### 2.3.2 Inference latency

Inference latency data was collected by simulating the process of executing the six models on images captured by the respective camera of each edge device. Studying inference response time for the smart dispenser were independent of the content captured in these images and corresponding predictions. Algorithm 1 describes our approach of computing average inference latency (10 images) over ten iterations for each model across the four edge devices. Tensorflow Lite (tflite-runtime) on Raspberry Pi SBCs and OpenMV’s Machine Learning module (ml) on the Arduino MCU devices were used to run inference. Python/MicroPython’s high-resolution timer time.time() was used to capture inference latency on all SBC and MCU devices. Model, images and result objects were deleted after computational use to clear volatile memory for subsequent iterations.

##### Algorithm 1

Simulating inference for smart bait dispenser

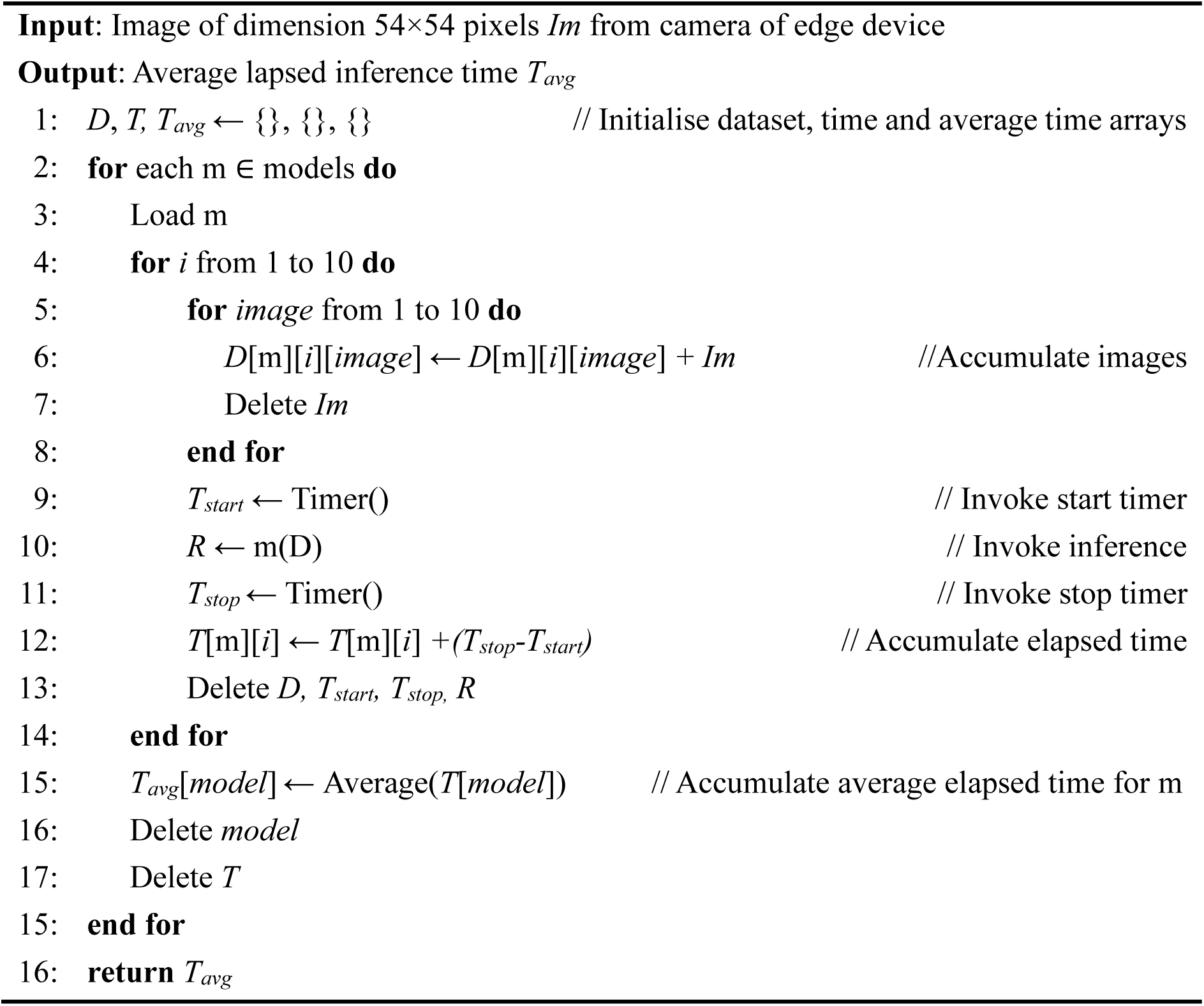

#### 2.3.3 Power consumption

We assessed the power consumption of each edge device in watt (W) across four operational states or modes-idle (or standby, not executing any task from program), capturing images and performing inference, deep sleep, and light sleep. Deep and light sleep are power-saving modes in MCUs that can be woken up with a timer or external interrupt to general-purpose input/output (GPIO) pins. Deep sleep minimises power consumption by powering down most internal modules, resulting in loss of volatile memory state. Light sleep retains volatile memory state by maintaining power to key internal modules. Thus, light sleep can resume operations from last operation executed before entering sleep mode. Since deep and light sleep modes are featured in only MCU, the “sleep” mode for SBCs was simulated with idle mode.

FNIRSI FNB38 universal serial bus meter (FNIRSI, 2020) was used to measure voltage in volt (V) and current drawn (I) in ampere (A), respectively, by the edge devices when running processes. The power consumption (P) was calculated by 𝑃 = 𝑉 × 𝐼. When the current draw was lower than one milliampere, a shunt resistor of 2.7 ohm was connected in series between ground and voltage input pins of the edge devices. The voltage drop was measured and used in calculation of current drawn by this formula: 𝐼 = 𝑉/𝑅.

## 3 RESULT

### 3.1 Image classification models

7452 training images and 1863 testing images were respectively used in training and assessing the binary classification models to identify Tasmanian devils from other species commonly found in Tasmania. Model sizes and test performance metrics across all full and quantised models is given in Table 2. CNN 1 was the smallest in size, followed by CNN2, MobileNetV3Small, MobileNetV2, ResNet50V2, and ResNet152V2, across all quantisation methods. As expected due to being simple models, CNN 1 and 2 were substantially smaller in size than pretrained models. A general dip in performance was observed in all quantised models compared to their respective original models; MobileNetV2 had a considerable improvement across all performance metrics (Figure 2). Quantising models typically reduces model performance, though opposite effects have been noted previously (example in (Van Baalen et al., 2023)). Custom trained CNN 1 achieved the highest accuracy, recall, and F1-score across all architectures and models. Although full sized CNN 1 had a higher precision than full sized CNN2, the latter achieved best precision of all the quantised counterparts.

**Table 2.**
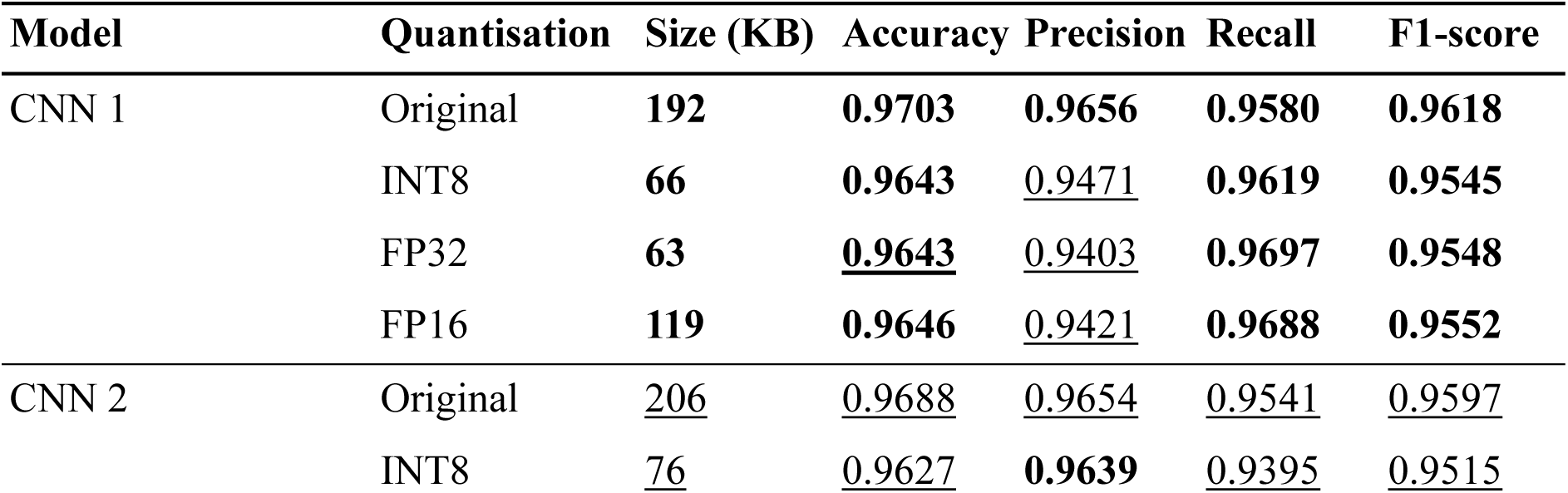
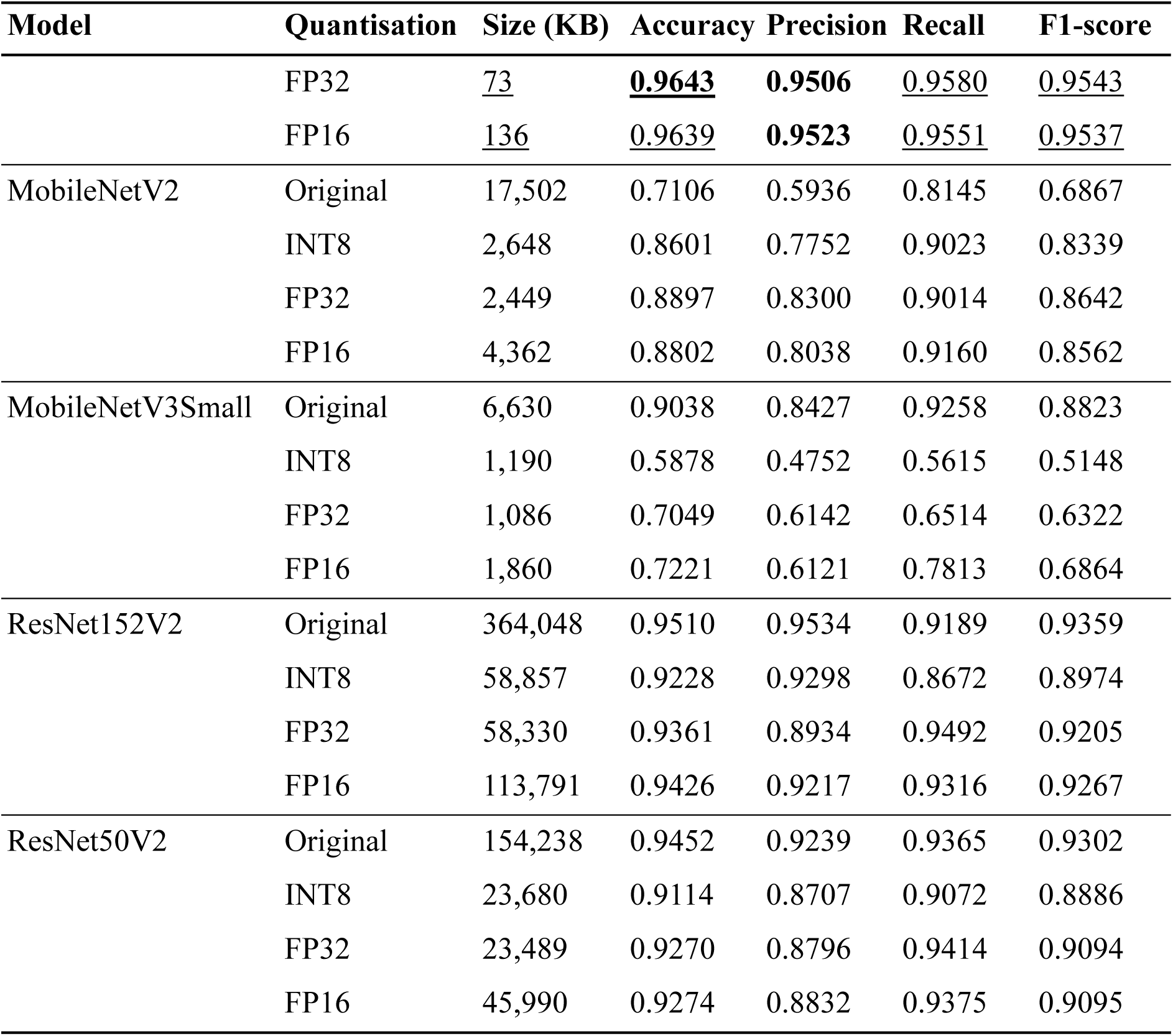
Comparison of size (in kilobyte) and test performance of original and post-training quantised models across metrics. Best model size (smallest) and metric scores (highest) per quantisation group are **bolded**, second-best are <u>underlined</u>.

Since our goal for training an image classification model is to use it in a smart bait dispenser for devils with potential of easy downstream adaptation for other target species, it is important that the model correctly identifies the target species, i.e., is high in sensitivity. This is because we would rather the dispenser deliver a bait to a non-target species occasionally than fail to drop a bait when a devil is present. Hence, recall (ratio of correctly predicted devils out of all devils in the dataset) should be considered as the primary metric of model evaluation. Additionally, as a complementary metric and indicator for holistic performance, we suggest that using F1-score to compare models. This metric provides a balance between precision and recall and is more insightful than accuracy when class imbalance is present in dataset, which is expected in real wildlife conservation scenarios. Since MCU boards can only support INT8 quantised models, we narrowed our focus on INT8 models only for head-to-head comparison across both MCU and SBC boards. INT8 quantised CNN 1 achieved the best results in both recall (96.19%) and F1-score (95.45%), making it the most suitable model for the smart bait dispenser based on test data evaluation.

### 3.2 Inference on edge devices

### 3.2.1 Latency

Observed inference latency is given in Table 3. The two SBCs— RPi 3B+ and RPi Z2W were able to run all the models, opposed to the MCU boards where Portenta could not run the two models of the ResNet family and Nicla Vision could not run any of the four pretrained models. The fastest average inference time of 0.069 seconds was observed on the Portenta running MobileNetV3Small, followed by 0.091 seconds on RPi Z2W running MobileNetV3Small, and 0.149 seconds on Nicla Vision running CNN 1. The slowest average inference time recorded was 2.836 seconds, when RPi 3B+ was running ResNet152V2. From these results, we consider inference times across all edge device to be quick enough for the bait dispenser. Hence, we must give a higher preference to INT8 performance metrics to identify the most suitable model for field deployment.

**Table 3.** Comparison of inference latency (in seconds) across different models. Lowest latencies under each model are **bolded**, second-lowest are <u>underlined</u>. N/A indicates data not available because model could not run on device.

| Device | CNN 1 | CNN 2 | MobileNet V2 | MobileNet V3Small | ResNet50 V2 | ResNet152 V2 |
| --- | --- | --- | --- | --- | --- | --- |
| ArduinoPro Nicla Vision | <u>0.149</u> | **0.150** | N/A | N/A | N/A | N/A |
| ArduinoPro Portenta H7 + Vision Shield (LoRa) | 0.253 | 0.255 | <u>0.158</u> | **0.069** | N/A | N/A |
| Raspberry Pi 3 Model B+ | 1.155 | 1.937 | 2.461 | 0.669 | <u>2.800</u> | <u>2.836</u> |
| Raspberry Pi Zero2 W | **0.124** | <u>0.187</u> | **0.251** | <u>0.091</u> | **0.257** | **0.262** |

### 3.2.2 Power consumption

Minimal power consumption will be critical for bait dispensers deployed in remote wilderness areas. Data on power consumption by the four edge devices across essential activity states are presented in Table 4. As expected, MCU boards utilise much less power than SBCs.

**Table 4.** Power Consumption (in watt) by selected edge devices across different operating modes. Lowest consumptions under each mode are **bolded**, second-lowest are <u>underlined</u>. N/A indicates data not available because device does not have sleep modes.

| Device | Idle | Running<br>Inference | Sleep mode |  |
| --- | --- | --- | --- | --- |
|  |  |  | Light Sleep | Deep Sleep |
| ArduinoPro Nicla Vision | <u>0.500</u> | <u>0.500</u> | <u>0.005</u> | **0.00187** |
| ArduinoPro Portenta H7 + Vision<br>Shield (LoRa) | **0.185** | **0.190** | **0.045** | <u>0.070</u> |
| Raspberry Pi 3 Model B+ | 3.400 | 3.650 | N/A | N/A |
| Raspberry Pi Zero2 W | 1.600 | 2.300 | N/A | N/A |

We observed that Portenta is ∼90% more efficient than Nicla Vision in both idle and inference modes, but the latter outperforms the former by 160% in light sleep and ∼190% in deep sleep. Upon discussion with Arduino developers (forum discussion in (kwagyeman, 2025)), we realised this is expected since Portenta is not optimised for low power. Between the two SBCs, RPi Z2W consumed less power than RPi 3B+ by 72% and 45.4% during idle and inference modes, respectively. Since SBCs do not offer sleep modes, the percentage difference of idle mode between the most power efficient SBC (RPi Z2W, 1.6 W) and the most efficient MCU board (Portenta, 0.185 W) is 158.5%. RPi Z2W in idle mode consumed 198.8% and 199.5% more power than Nicla Vision’s light sleep (0.005 W) and deep sleep (0.00187 W) modes, respectively.

## 4 DISCUSSION

Beyond the immediate goal of Tasmanian devil vaccination, this study introduces a broadly adaptable method for management of species and ecosystems. The use of a camera-based edge device with onboard deep learning offers a pathway for automated wildlife monitoring, invasive species control, and targeted feeding programs. By integrating field-ready computer vision with local decision processing, this work bridges technological development and applied conservation practice.

In this study, we compared four edge devices to assess the feasibility of using them in a smart bait dispenser for vaccination of Tasmanian devils against DFTD. We short listed two generic and four pretrained CNN-based image classification models to deploy on the considered devices. CNN1 maintained stable performance (recall 96.2 ± 0.3%, F1-score 95.5 ± 0.2%), indicating consistent classification behaviour. From an ecological perspective, this level of sensitivity is desirable, as occasional false positives (i.e., dispensing a bait to a non-target species) are less critical than missing a target individual that should receive a vaccine. We conclude CNN 1 to be the best performing model to classify devil from non-devil species based on test data evaluation on metrics recall and F1-score, and average inference latency across the four devices.

The image classification approach allows for rapidly adapting the models for alternate target species. Existing camera trap images on any target species like racoons and striped skunks (*Mephitis mephitis*) in North America can be leveraged for applications like vaccination. An eastern quoll translocation project is underway to help restore genetic diversity in declining populations in Tasmania (Hamer et al., 2023). Supplemental feeding has been used to provide food while the quolls navigate the new site. The smart bait dispenser could be repurposed to deliver the supplemental food only to quolls. In this case, our dataset already contains eastern quoll images, so reusing the species recognition models only requires defining eastern quoll as the target species and retraining.

Vaccination campaigns are typically multiday field projects and hence the smart bait dispenser must be power conscious. ArduinoPro Portenta H7 Vision Shield (LoRa) yielded quick inference results and the lowest power consumption during idle state and inference. Having a sleep mode on the dispenser is essential because the system will remain inactive for majority of time in wait until a motion sensor triggers the process to start. An efficient sleep mode will extend the battery life of the dispenser and keep it running in field for days with low to no battery maintenance. As mentioned earlier, Portenta is not optimised for sleep and SBCs RPi 3B+ and RPi Z2W do not have sleep modes. Nicla Vision yielded the lowest deep and light sleep power consumption of 0.002 W and 0.005 W respectively. However, a low-end MCU like Atmel ATtiny85 can attain a much lower deep and light sleep power consumption of up to 6 μW and 30 μW respectively (Atmel, 2013). Hence, ATtiny85 in sleep mode can be used as a low power switch that turns on the primary edge device upon motion. Using this method with RPi 3B+ and RPi Z2W is not suitable for the dispenser because these SBCs have a substantially longer boot-up times than MCUs.

Between the two MCU boards in this study, the 28-pin Portenta has prototyping advantages over the 19-pin Nicla Vision. Portenta’s broader input/output capabilities support more extensive external integrations with the bait dispenser. Although both the devices have dual-core processors, Portenta has a larger memory of 8 megabytes (MB) of synchronous dynamic random-access memory and 16 MB flash, whereas Nicla Vision only has 1 MB random-access memory and 2 MB flash. Additionally, Portenta’s storage can be conveniently expanded via a memory card using the Vision Shield, streamlining data logging processes. In contrast, Nicla Vision requires a separate compatible memory card reader module to achieve similar functionality.

The GC2145 (2 megapixel) camera of Nicla Vision is superior to HM01B0 (320×240 pixels) camera of Portenta Vision Shield. However, GC2145 includes infrared (IR) filter by default, making it unsuitable to capture images during dark under infrared lights unless removed manually. HM01B0 lacks IR filter and hence can be used in the dispenser. Newer iterations of Portenta Vision Shield are manufactured with upgraded HM0360 camera module (640×480 resolution) that has IR filter. A no-IR variant of HM0360 can be sourced directly from Arducam, the official manufacturer.

The long-range (LoRa) radio communication variant of Portenta Vision Shield can enable field scientists to remotely monitor the bait dispenser via long-range wide area network (LoRaWAN). To achieve this with other edge devices considered in this study, we would have to use an external LoRa module. LoRaWAN is an attractive functionality addition to the dispenser as it can be used to display live updates on motion triggers, species prediction, and number of baits dispensed on an online dashboard. Applications of LoRaWAN are found in agriculture and animal monitoring ((Boo et al., 2023; Haxhibeqiri et al., 2018; Hunan HKT Technology Co.) and reviewed in (Schulthess et al., 2023)) due to key features like security, low power consumption, ease of deployment and network coverage of over 15 km in clear line of sight in rural areas (Petajajarvi et al., 2015).

Altogether, we found that simple traditional CNNs was best suited for species classification in a camera-based smart bait dispenser for vaccination of Tasmanian devils. We conclude that ArduinoPro Portenta H7 Vision Shield (LoRa) is the most competent device for this edge AI application. Additionally, we suggest using Atmel ATtiny85 microcontroller at achieve sleep mode on Portenta for multi-day field application.

Future studies will need to validate prototypes of the proposed smart bait dispenser. Field trials with the prototype will evaluate deployment strategy, battery endurance, and resistance to weather exposure. Integration of the dispenser with LoRaWAN will allow real-time monitoring of activity and bait uptake. The framework can be extended to other conservation contexts such as population monitoring and supplemental feeding in eastern quolls to highlight its versatility as a general ecological tool.

## Acknowledgements

We thank the Department of Natural Resources and Environment Tasmania for providing trail camera images. This study was funded by the Australian Research Council Linkage Projects grant LP210301148 awarded to Andrew S. Flies.

## Data availability

Data will be made available upon request.

## Conflict of interest statement

The authors declare no competing interests.

## Author contribution

Prithul Chaturvedi, Andrew S. Flies, Soyeon Caren Han and William M. Connelly conceptualised the approach of using image classification for target-specific bait delivery. Andrew S. Flies enabled the procurement of trail camera images for model development. Prithul Chaturvedi, Soyeon Caren Han and William M. Connelly developed the models and conducted experiments on the edge devices. Prithul Chaturvedi led the writing of the manuscript. All authors contributed critically to the drafts and gave final approval for publication.

## Notes

### Competing Interest Statement

The authors have declared no competing interest.

### Summary of Updates

Author updated; Citation formatting revised; Section 4 updated to clarify.

